# Optimising a Simple Fully Convolutional Network (SFCN) for accurate brain age prediction in the PAC 2019 challenge

**DOI:** 10.1101/2020.11.10.376970

**Authors:** Weikang Gong, Christian F. Beckmann, Andrea Vedaldi, Stephen M. Smith, Han Peng

**Author notes:** Correspondence: Han Peng.

## Abstract

Brain age prediction from brain MRI scans not only helps improve brain ageing modelling generally, but also provides benchmarks for predictive analysis methods. Brain-age delta, which is the difference between a subject’s predicted age and true age, has become a meaningful biomarker for the health of the brain. Here, we report the details of our brain age prediction models and results in the Predictive Analysis Challenge 2019. The aim of the challenge was to use T1-weighted brain MRIs to predict a subject’s age in multicentre datasets. We apply a lightweight deep convolutional neural network architecture, Simple Fully Convolutional Neural Network (SFCN), and combined several techniques including data augmentation, transfer learning, model ensemble, and bias correction for brain age prediction. The model achieved first places in both of the two objectives in the PAC 2019 brain age prediction challenge: Mean absolute error (MAE) = 2.90 years without bias removal, and MAE = 2.95 years with bias removal.

## 1 Introduction

Predictive analysis with data-driven machine learning algorithms brings huge promise in neuroimaging and neuroscience research. Predictive analysis can not only help disease diagnosis, such as Alzheimer’s (Liu et al., 2018), Autism (Thomas et al., 2020), ADHD (Zou et al., 2017) and schizophrenia (Zeng et al., 2018), but also helps in formulating new hypotheses (Shmueli, 2010) and identifying new biomarkers (Rosenberg et al., 2018). Yet, the predictive analysis paradigm brings new challenges. First, a fair way to compare predictive analysis models is needed. In predictive analysis, it is common practice to build models in a training set, and then apply the models to a test set (Bzdok et al., 2020; Scheinost et al., 2019). It is important that no test data is used for model training or hyperparameter tuning (e.g. learning rate for gradient decent optimisations, number of layers in convnets) and to report the result objectively (LeCun et al., 2015) and avoid accidental data leakage (Lanka et al., 2019). Second, data is usually scarce for many diseases so that training a large deep learning model in such modest datasets is still hard (Raghu et al., 2019).

Brain ageing study is a recent example of the predictive analysis paradigm (Brown et al., 2012; Cole et al., 2018, 2017; Cole and Franke, 2017; Dosenbach et al., 2010; Franke et al., 2010; Levakov et al., 2020; Neeb et al., 2006). Studies showed that individuals’ chronological age can be predicted accurately from brain MRI scans (Cole et al., 2017). Brain age delta, the difference of a subject’s predicted (brain) age and chronological age, is linked with a variety of biological factors within the healthy population (Smith et al., 2020b), and group differences can be found in disease populations (Cole et al., 2019; Kaufmann et al., 2019). Yet, accurate prediction of a subject’s age in healthy population is still a challenging task.

To tackle these challenges, a benchmarking platform is needed to objectively evaluate the models and strategies. Competitions have been seen in the field of computer vision (e.g. ImageNet (Russakovsky et al., 2015)) and proved to be a valuable vehicle for pushing AI technology (LeCun et al., 2015). In the field of neuroimaging, the Predictive Analysis Challenge (PAC) 2019 for brain age prediction^1^ provides such opportunities for participants to train machine learning methods, and then objectively evaluate the models in a test dataset whose labels are hidden from the participants. PAC 2019 sets two objectives for brain age predictions: (1) to achieve the most accurate age prediction from brain structural MRI scans, and (2) to achieve the best accuracy while keeping the correlation between the prediction error and the ground truth age sufficiently small.

Our team ‘BrainAgeDifference’ achieved the first places in both two objectives among 79 participating teams. Our method is largely based on our previous work (Peng et al., 2019), with adaptations made for the challenge. In this report, we will provide a detailed description of our methods for PAC 2019, including the lightweight deep convnet architecture - Simple Fully Convolutional Neural Network (SFCN), and the combined techniques including data augmentation, transfer learning, model ensemble, and bias correction. We find that the lightweight model, which has achieved the state-of-the-art results in UK Biobank, works well in the multi-centre PAC 2019 dataset with a slightly adaptation in hyperparameters. SFCN pretrained on UK Biobank data achieves better single model performance than random initialized models in the PAC 2019 dataset. In addition, model ensemble with different T1-image derived maps, and different initializations, and training/validation data splits are important to achieve the best performance for the competition.

## 2 Datasets and Preprocessing

### 2.1 PAC 2019

The Predictive Analytic Challenge (PAC) 2019 was to predict age from brain MRI scans. The goal of the challenge includes two parts: (1) to achieve the most accurate age prediction, as measured by mean absolute error (MAE), and (2) to achieve the best MAE while keeping the Spearman correlation r-value between the prediction error (brain age delta) and the actual age below 0.1 (|*r*| <0.1). The dataset consists of both label-known training/validation dataset (2638 subjects in total) and a ‘true’ test set of 660 subjects whose labels are unknown to the competition participants. The participants had a one-time opportunity to upload their predictions in the test set to the competition server for each objective, and the MAE and the Spearman’s r-value were evaluated automatically. The subjects are from 17 different sites. Most of the data is based on (Cole et al., 2017) and a few new sites were added by the organisers. The training set and the test set have the same age and site distribution.

PAC 2019 organizers provide three version of MRI data: (a) raw T1 brain MRI scans, (b) white matter volume segmentation (WM) and (c) grey matter volume (GM) segmentation derived from T1 data. We use all three versions to develop deep learning models. We further preprocess the raw T1 images using FSL (Smith et al., 2004) (command fsl_anat) to derive two different pseudo-modalities: one is brain linearly registered to standard 1mm MNI space (by FLIRT), and the other is brain non-linearly registered to standard 1mm MNI space (by FNIRT). We use all the four pseudo-modalities to develop the convnet models. WM and GM segmentations are in 1.5mm MNI space as provided by the PAC 2019 organisers, and the preprocessing pipeline is described in (Cole and Franke, 2017).

For linearly and non-linearly registered modalities, the input images are cropped to retain the central 160×192×160 voxels, which is the same as what we had done with UK Biobank data. The WM and GM modalities are cropped in the central 96×128×96 voxels.

### 2.2 UK Biobank

UK Biobank brain imaging data consists of multimodal brain scans from a predominantly healthy cohort (Miller et al., 2016). Currently (year 2020) there are about 40,000 subjects released for research, and the number will eventually reach 100,000 (Smith et al., 2020a). In our previous study, we reported SFCN trained and tested on the initial 14,503 structural MRI brain images (Peng et al., 2019), and released the pretrained model in a GitHub repository (https://github.com/ha-ha-ha-han/UKBiobank_deep_pretrain). In this study, we mainly focus on optimising pipelines and models for PAC 2019, and most of the models are initialised randomly and then trained with the PAC 2019 data unless otherwise stated. To apply transfer learning, we also use 5698 UK Biobank T1 images to pretrain a model, and then use the trained weights as initialisations for finetuning five models in the PAC 2019 dataset (see details in the section Experiments and Results – Transfer Learning).

The UK Biobank preprocessing pipeline can be found in (Alfaro-Almagro et al., 2018), and the UKB data release includes preprocessed data, so that researchers do not need to re-run the preprocessing pipeline. Models are trained/validated/tested separately. The inputs are in 1mm MNI space, cropped for the central 160×192×160 voxels to reduce GPU memory required.

### 2.3 Difference between UK Biobank and PAC 2019

UK Biobank and PAC 2019 datasets differ in age distribution and number of subjects. A summary of the statistics of both datasets (mean and standard deviation of age distribution, and number of subjects) is shown in Table 1 and visualised in Figure 1. The PAC 2019 dataset has a significantly smaller number of subjects and larger age range. Moreover, PAC 2019 contains multisite data with different data quality and scanner configurations. All these factors make the prediction task more difficult in PAC 2019 than UK Biobank.

**Table 1.** **Difference in age distribution between PAC 2019 used in this study and UK Biobank dataset used in (Peng et al., 2019)**.

| Dataset | Age Range (yrs) | Age (yrs) Mean±STD | Number of Subject | | Number of Site |
| --- | --- | --- | --- | --- | --- |
|  |  |  | Training/Validation/Test | Total |  |
| UK Biobank | 44 - 80 | 62.7±7.5 | 5698 / 518 / - | 6216 | 2 |
| PAC 2019 | 17 - 90 | 35.9±16.2 | 2198 / 440 / 660 | 2638 with label + 660 without label | 17 |

**Figure 1.**
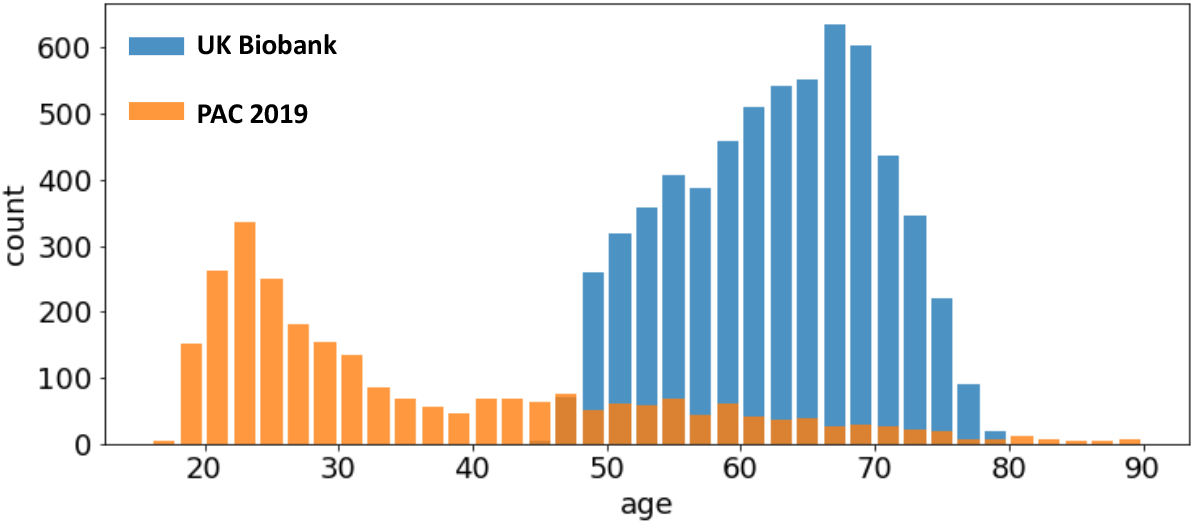
Age distribution of different datasets. The UK Biobank (blue bars) and the PAC 2019 (orange bars) differ in age range and number of subjects.

Note that the test set labels are not available to the participants in the PAC 2019 challenge. This setup of a ‘true’ test set prevents the competition participants from the risk of accidental data leakage. During the competition, the prediction results were allowed to be uploaded only once, and then the performance metric was evaluated automatically. Therefore, no hyperparameter adjustment could be made for the testing process to elaboratively overfit the test set. In summary, we believe the results in the test set are an objective measurement of model performance in an unknown dataset with a similar age and site distribution.

## 3 Method

### 3.1 Model

The backbone of our method is the lightweight fully convolutional neural network architecture, Simple Fully Convolutional Neural Network (SFCN), that we proposed in (Peng et al., 2019). We briefly summarise the key aspects of the model and the adjustment for PAC 2019 here.

The SFCN model architecture is shown in Figure 2 (reproduced from the original work by (Peng et al., 2019)). The model consists of seven convolution blocks. Each of the first five blocks consist of a 3×3×3 3D convolution layer, a batch normalisation layer, a max pooling layer, and a ReLU activation layer. The key facet of this architecture is that the model downsamples the input every time after a convolution layer. As a result, the spatial dimension is reduced quickly as the layer goes deeper, and it takes only five blocks to reduce the input data size from 160×192×160 to 5×6×5 (voxels). This simple design saves GPU memory and reduced the number trainable weights. The sixth block is similar but without a max pooling layer and uses a 1×1×1 3D convolution layer to increase non-linearity without changing feature map spatial dimensions. The resulting 5×6×5 feature map is pooled by an average pooling layer and then projected to the output layer with a linear transformation (i.e. fully connected layer). For convenience of implementation, the fully connected layer is also treated as an 1×1×1 Conv3D in a 1×1×1 input ‘feature map’.

**Figure 2.**
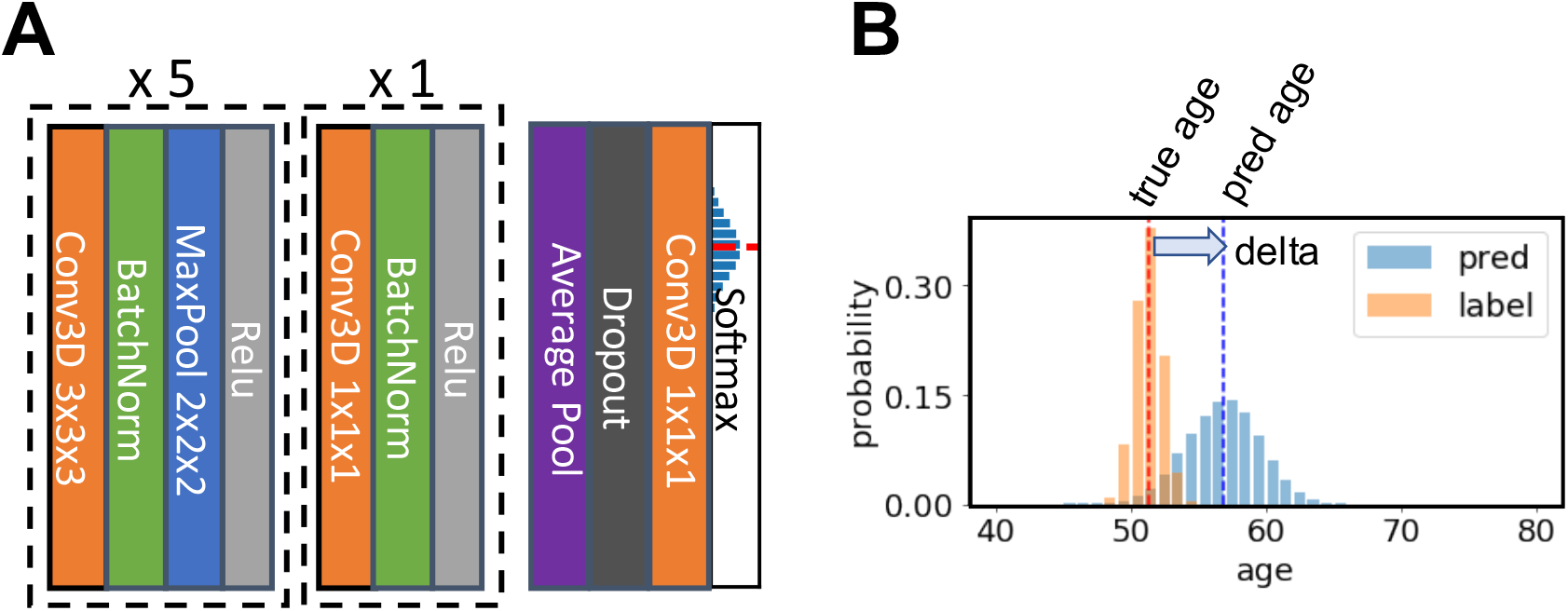
Illustration of the core network for the Simple Fully Convolutional Neural Network (SFCN) model. A) SFCN model architecture. B) An example of soft labels and output probabilities. The figure is reproduced from (Peng et al., 2019) under CC-BY-NC-ND 4.0.

The input size is 160×192×160 voxels for both T1 non-linearly registered brains and linearly registered brains, and 96×128×96 voxels for both WM and GM for PAC 2019. Note that the model is fully convolutional; therefore it can take different input sizes without modifying the architecture. The feature map size before the average pooling layer in the final block is 5×6×5 for the input size 160×192×160, and 3×4×3 for the input size 96×128×96.

### 3.2 Model Output and Loss function

We treat the regression as a soft classification problem. In this set-up, the label of the age is not treated as a single number, but a discretized Gaussian probability distribution centred at the true age. The output of the model is also a probability distribution. Kullback-Leibler divergence is used to measure the similarity between the two probabilities.

The output is 40 digits standing for 40 age bins for the UK Biobank data. Each age bin covers a 1-year range. The number of age bins is 38 for trained-from-scratch models for PAC 2019, each of which covers a 2-year range. The sigma of the Gaussian distribution for the labels is set to be the size of one age bin (i.e. 1 year for UK Biobank and 2 years for PAC 2019). The final age prediction is the average of all the age bins weighted by the output probability. For models pretrained in UK Biobank and finetuned in PAC 2019, the number of output age bins is set to 40 to reduce coding effort (although the bins stand for different age ranges).

### 3.3 Hyper parameter, optimiser choice and training

Hyper parameters are tuned with the validation set. We also evaluate different optimizers, namely, ADAM and SGD. In UK Biobank we find ADAM easily overfits the model and thus performs worse than SGD (Peng et al., 2019). However, in PAC 2019, we find that ADAM, although it overfits more than SGD (as measured by the val-train gap in Figure 3), performs slightly better than SGD in the validation set. Also, ADAM is observed to be more stable during the training process for the PAC 2019 dataset (as shown in Figure 3), so that we use ADAM for PAC 2019 for the rest of our experiments.

**Figure 3.**
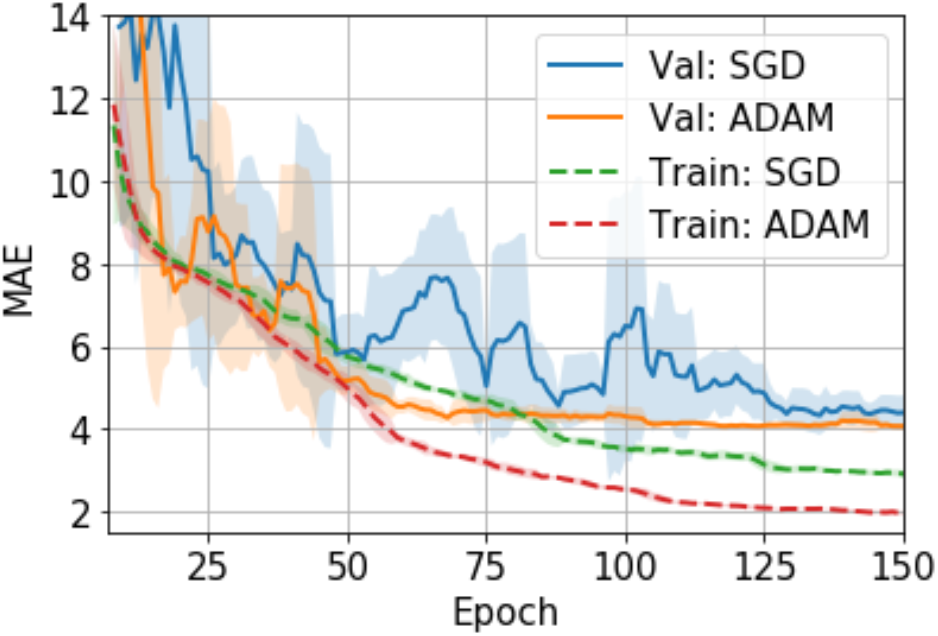
Training curves for the SGD and ADAM optimisers in PAC 2019 data. The curves are smoothed with a 7-step averaging window. The shading areas show the standard deviation within the window.

The validation set is used to evaluate model performance after every epoch (i.e. one iteration through the full dataset) in the training set, and the model weights for the best validation performance within 150 epochs are chosen for testing. Data augmentation and weight regularization are important to achieve the best prediction accuracy and to reduce overfitting. We use the same augmentation and regularization strategy as specified in detail in (Peng et al., 2019) for all experiments reported in this work: voxel shifting, mirroring and dropout.

## 4 Experiments and Results

To achieve accurate brain age prediction, we use several techniques in the competition setup besides the lightweight SFCN model, the regularization and the data augmentations. For a single model, we applied transfer learning to boost the single model prediction accuracy. We also train multiple models using different (pseudo-)modalities to form an ensemble for better performance. As summarised in Table 2, we find that the best ensemble uses all the modalities. While transfer learning stably achieves better single-model performance, only 5 out of 45 models in the final ensemble are transferred from UK Biobank, due to the limit of time and computational power. The details of the experiments and the results are described below.

**Table 2.** Performance of model ensembles with different pseudo modalities in PAC 2019. 5 models are initialized with pretrained weights and then finetuned with linearly registered brains. For all other experiments, 10 models are trained from scratch for each modality and used to predict brain age individually. The mean and the standard deviation of the single model performances are computed within each modality.

| Modality | Performance |  |  |  |
| --- | --- | --- | --- | --- |
|  | Single Model |  | Ensemble |  |
|  | MAE (yrs) | r value | MAE (yrs) | r value |
| Raw, linearly registered,<br>Pretrained with UK<br>Biobank x 5 | 3.69±0.08 | 0.946±0.006 | 3.22 | 0.960 |
| Raw, linearly registered<br>x 10 | 3.91±0.13 | 0.935±0.007 | 3.48 | 0.951 |
| Raw, non-linearly<br>registered x 10 | 3.89±0.16 | 0.937±0.006 | 3.40 | 0.957 |
| Grey matter<br>x 10 | 3.93±0.13 | 0.948±0.003 | 3.54 | 0.957 |
| White matter<br>x 10 | 4.19±0.09 | 0.937±0.003 | 3.74 | 0.951 |
| All 45 models | 3.95±0.19 | 0.940±0.007 | 2.98 | 0.971 |

### 4.1 Transfer learning

To test how pretraining in the large UK Biobank dataset can help smaller datasets such as PAC 2019, we compare the performance of models that are pretrained-and-finetuned and those trained-from-scratch using the PAC 2019 data only.

The finetuning process and all the hyperparameters are the same with the trained-from-scratch ones except for the initialisation of model weights. For the pretraining, an SFCN model is trained with 5698 UK Biobank subjects using the methods specified in (Peng et al., 2019) and achieving validation MAE = 2.20 yrs in UK Biobank dataset. This MAE is slightly worse than the reported value due to the smaller training dataset size we use. The trained weights are then used to initialise models that are finetuned with the PAC 2019 dataset. There are five models initialised with the same weights, and then trained with different train-validation split under a five-fold cross validation scheme using the PAC 2019 training data. as shown in Figure 4, the five finetuned models achieve a mean MAE of 3.69±0.19 yrs (mean±STD), which is 0.22 years better than the randomly initialised models (MAE = 3.91±0.13 yrs, mean±STD). The pretrained models also converge faster. This result shows that initialising models with pretrained weights from UK Biobank can help achieve better performance in small datasets, even using a naïve finetuning protocol.

**Figure 4.**
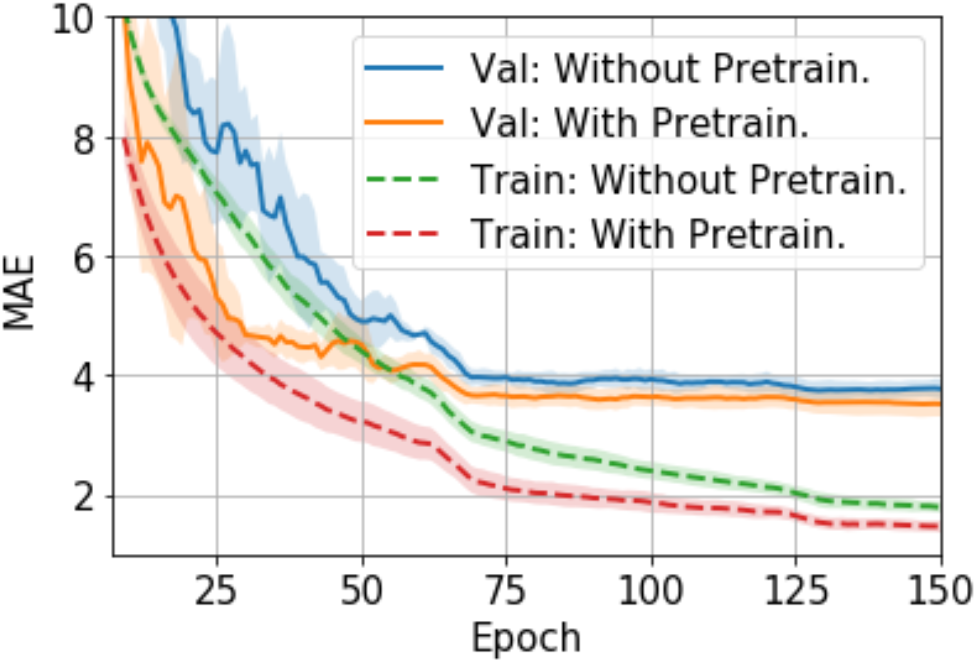
Training curves for transfer learning. The curves are averaged by five models trained with five-fold cross-validation splitting, and then smoothed with a 7-step averaging window. The shading areas show the standard deviation within the window.

### 4.2 Performance of different (pseudo-)modalities and model ensembles

Different T1-derived data contain distinct information regarding brain ageing. We find that averaging predictions with different pseudo-modalities (outputs from distinct pre-processing approaches applied to the same original input data modality, here T1) is an effective method to utilise the independent information to achieve the overall best ensemble performance. We train and test 10 models (from scratch, no pretraining) in each pseudo-modality, namely, T1 data linearly registered to the MNI space (Lin), raw T1 data nonlinearly registered to the MNI space (NonLin), segmented grey matter (GM) and white matter (WM) volumes. Lin and NonLin modalities are preprocessed by us, and GM and WM are provided by the organiser. Models are randomly initialized (with different random seeds). As shown in Table 2, models trained with Lin, NonLin and GM achieve comparable MAEs ranging from 3.89 to 3.93 years, which are all better than the MAE for WM (4.19 years), and is in accordance with our previous findings (Peng et al., 2019).

We show in our previous work (Peng et al., 2019) that, even though with comparable MAEs, brain-PADs contain different information from different pseudo-modalities. This result is consolidated in the PAC 2019 dataset using the left-out validation set (not used in cross-validation) in Figure 5. Models with the same modalities show higher correlation for the brain-PAD prediction.

**Figure 5.**
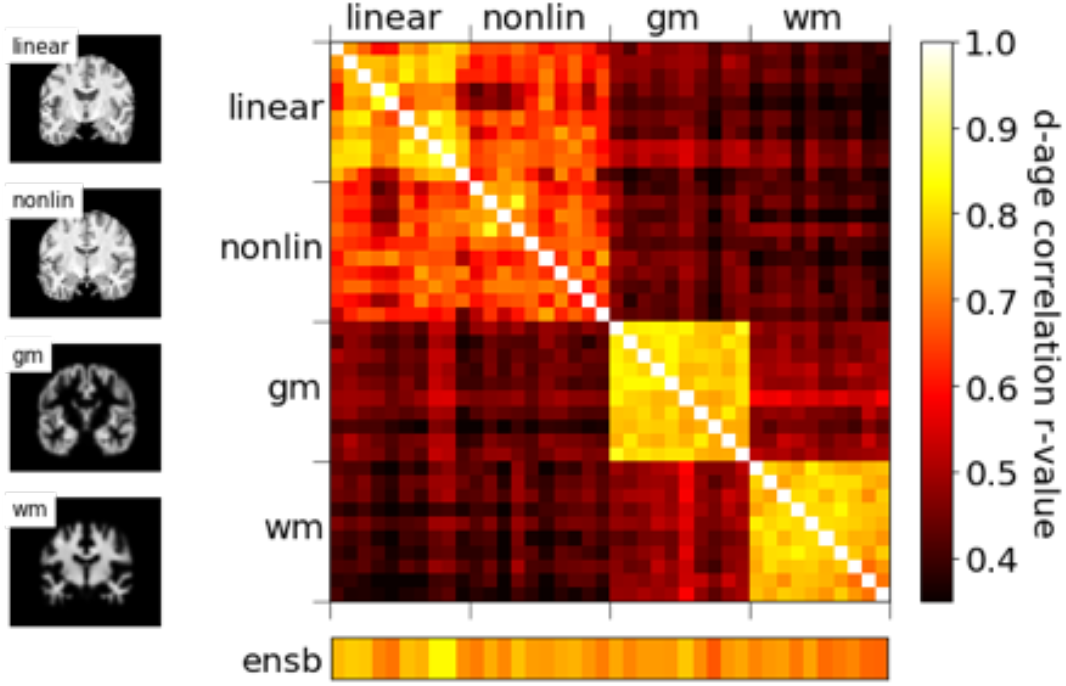
**Correlations of predicted brain age difference (d-age) between different models, showing similar results as (Peng et al., 2019)**.

To achieve the best performance in the challenge, we use all four pseudo-modalities to form an ensemble. For every pseudo-modality, there are 10 models initialised randomly and trained separately with different train/validation splits. For the Lin modality, 5 additional models are pretrained in UK Biobank and finetuned in PAC 2019, as previously mentioned, adding up to 45 models in total. All models are trained separately, and make predictions independently. For every subject, mean and standard deviation (STD) are computed for the 45 age predictions, and the predictions deviating more than *λ*-STD from the mean are treated as outliers (*λ* is a coefficient of our choice), and the final prediction is the new average of the rest predictions. *λ* is set to be 1.1 to optimise the performance in the left-out validation set, which makes the ensemble performance slightly biased towards this ‘validation’ set. This strategy achieves MAE = 2.98 yrs in the left-out validation set and MAE = 2.90 yrs in the test set, as shown in Table 3. Our result in the test set ranks the first for the first goal of PAC 2019 (best MAE), and is 0.18/0.42 years better than the second/third place (MAE: Ours = 2.904 yrs; Second Place = 3.086 yrs; Third Place = 3.328 yrs).

**Table 3.** Bias correction results.

| Model | Performance |  | Performance with Bias Correction |  |
| --- | --- | --- | --- | --- |
|  | MAE (years) | Spearman Correlation<br>d-age vs age | MAE (years) | Spearman Correlation<br>d-age vs age |
| 45 Model Ensemble<br>(Left-out validation set) | 2.98 | -0.44 | 3.01 | -0.06 |
| 45 Model Ensemble (PAC Test Set) | 2.90 | -0.39 | 2.95 | -0.03 |

In our previous work (Peng et al., 2019), we showed that independent predictions are important to form a good ensemble. Here, we further show that a sufficiently large number of models is also important for good ensemble performance. To demonstrate this, we explore the ensemble performance with different number of models, as summarised in Figure 6. Ensembles are randomly formed using some of the 45 trained models (replacement allowed) and predictions are made using the mean without excluding outliers. As the number of models increases, the MAE decreases and finally saturate. A power law can be fitted to empirically describe the quantitative relationship between the size of ensemble and the MAE, as shown in Figure 6B. A ‘critical point’ of MAE of 3.07 yrs is estimated, and can be interpreted as the ideal MAE if we can increase the number of models to infinity. This empirical observation suggests that simply increasing ensemble size will result in only limited performance gain.

**Figure 6.**
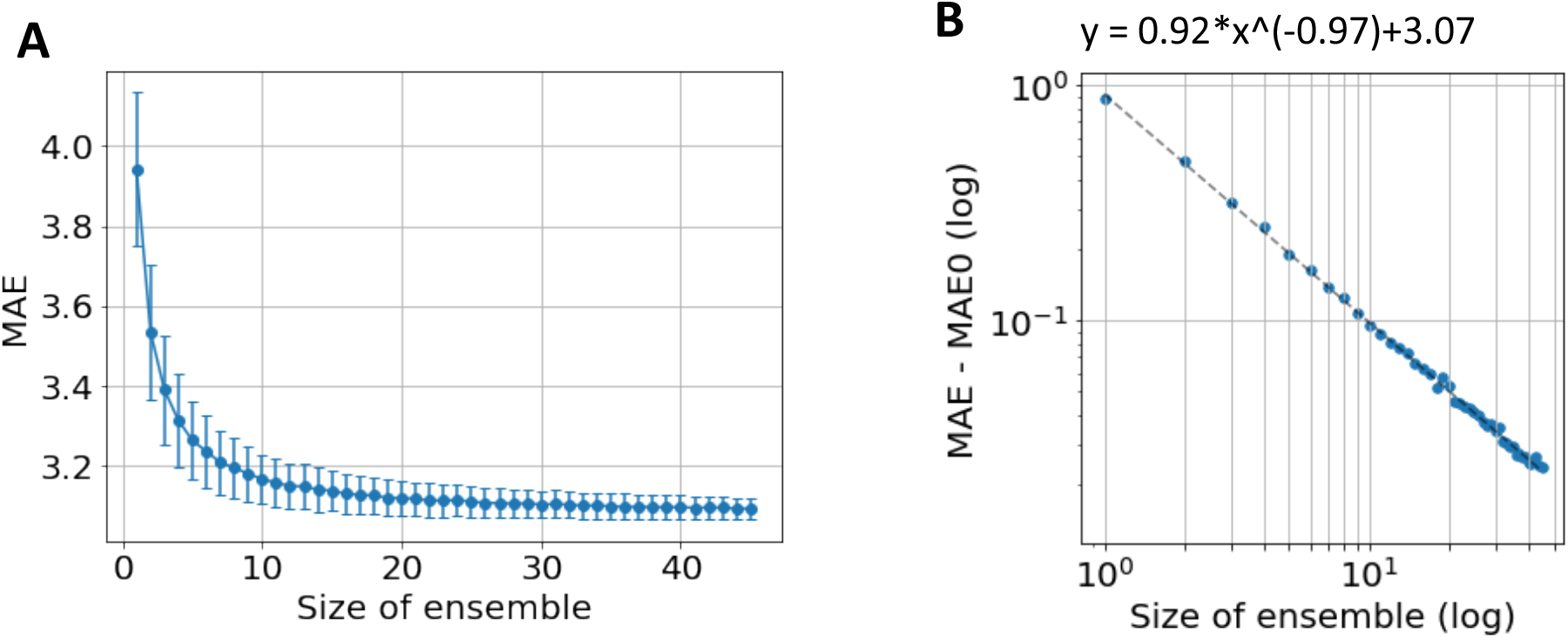
Ensemble performance with different number of models. **A)** Average performance in MAE with different number of models used by ensemble. The mean and standard deviation come from 1000-time bootstraps. **B)** The fitted line of a power law. *MAE*_0_ is the critical point if an infinite number of models are used to form the ensemble.

The ‘critical’ MAE is worse than the actual MAE we get from the all the models. This is because the bootstrap process allows replacement, i.e. the same model is allowed to be selected more than once, which reduces the independent information gathered from the ensemble.

### 4.3 Bias correction

We follow (Smith et al., 2019) and (Peng et al., 2019) to fit a straight line between the predicted brain-PAD and the ground truth age in the left-out validation set, and then apply the fitted parameters (slope and intercept) to bias-correct predictions in the test set whose labels are unknown. We correct the bias for the ensemble predictions rather than for every single model.

For the validation set, this linear regression method reduces the Spearman’s r-value (between delta and age) from -0.44 to -0.06 with a small increase (0.03 years) in the MAE. The generalization to the test set reduces the Spearman’s r-value from -0.39 to 0.03, with a small increase of 0.05 years in the MAE (from MAE = 2.90 to MAE = 2.95). This result is summarised in Table 3.

The result in the test set achieves the first place for the second goal of the competition (smallest MAE with sufficiently small Spearman’s r-value between brain-PAD and the true age), and it leads by a large margin (MAE: Ours = 2.950 yrs; Second Place = 3.799 yrs; Third Place = 3.924 yrs).

## 5 Discussion and conclusion

We note that different datasets may require distinct hyperparameters and optimisers for optimal performance for a deep learning algorithm. For example, we showed in our previous study that ADAM easily overfits the model and thus performs worse than SGD in UK Biobank data (Peng et al., 2019). In this study, we find ADAM works comparable or even slightly better than SGD in PAC 2019 validation data. We have not fully explored the mechanism behind this empirical difference. One can assume that PAC 2019 is a more difficult dataset for deep learning models to optimize, due to the multi-site origin and inhomogeneous data quality, and this may be the reason why ADAM performs better in PAC 2019; it has been shown to be a more powerful optimizer for other problems (Kingma and Ba, 2014). For future studies, it may be beneficial to explore and choose different optimisers for different datasets even for similar tasks.

Despite additional hyperparameter tuning, we have shown that the SFCN method together with the data augmentation and model regularisation methods are generalisable outside the UK Biobank dataset. However, this ‘generalisability’ requires retraining or finetuning in the targeting dataset, and may not be feasible for smaller datasets (e.g. a dataset with 100-subject). Also, although PAC 2019 provides a true measurement for generalisability of models to unseen data (because the test set labels are hidden from the participants), this does not guarantee the generalisability to unseen scanning site (because the test set follows the same site and age distribution as the training set). For applications requiring site generalisability, see recent work aiming to address this specific issue (Dinsdale et al., 2020).

Finally, we need to point out that our choice of hyperparameters, transfer learning and the naïve ensemble strategy may not be optimal, due to the limit of time and computation power in the competition setup.

To conclude, we have applied the lightweight convnet - SFCN model, data augmentation, regularisation, and bias correction techniques proposed in (Peng et al., 2019) to PAC 2019 challenge and achieved leading results. Besides initialising models randomly, we have shown that initialising weights pretrained in UK Biobank achieve better single-model results for the PAC 2019 dataset (after retraining/finetuning). For ensembles with multiple models, we have shown that the best ensemble comes from a large number of models taking the input of different pseudo-modalities.

## 6 Conflict of Interest

*The authors declare that the research was conducted in the absence of any commercial or financial relationships that could be construed as a potential conflict of interest*.

### 7 Author Contributions

**Weikang Gong:** Conceptualization, Methodology, Software, Writing - Review & Editing. **Christian F. Beckmann:** Conceptualization, Writing - Review & Editing, Methodology, Funding acquisition, Supervision. **Andrea Vedaldi:** Conceptualization, Writing - Review & Editing, Methodology, Funding acquisition, Supervision. **Stephen M. Smith:** Conceptualization, Writing - Review & Editing, Methodology, Funding acquisition, Supervision. **Han Peng:** Conceptualization, Methodology, Software, Writing-Original draft preparation, Writing - Review & Editing, (Co-)Supervision.

## 9 Acknowledgments

This research has been conducted in part using the UK Biobank Resource under Application 8107. We are grateful to UK Biobank for making the data available, and to all UK Biobank study participants, who generously donated their time to make this resource possible. Computation used the Oxford Biomedical Research Computing (BMRC) facility, a joint development between the Wellcome Centre for Human Genetics and the Big Data Institute supported by Health Data Research UK and the NIHR Oxford Biomedical Research Centre.

## 10 Data Availability Statement

The PAC 2019 dataset consists of several public available datasets and a few datasets provided by the organiser. Interested researchers can apply for the access to the public available datasets as specified in (Cole et al., 2017) and need to contact the PAC 2019 organisers for the rest of the sites. The UK Biobank dataset is accessible upon applications via the website: https://www.ukbiobank.ac.uk/

## Footnotes

1 https://web.archive.org/web/20200214101600/https://www.photon-ai.com/pac2019

## References

Alfaro-Almagro, F., Jenkinson, M., Bangerter, N.K., Andersson, J.L.R., Griffanti, L., Douaud, G., Sotiropoulos, S.N., Jbabdi, S., Hernandez-Fernandez, M., Vallee, E., Vidaurre, D., Webster, M., McCarthy, P., Rorden, C., Daducci, A., Alexander, D.C., Zhang, H., Dragonu, I., Matthews, P.M., Miller, K.L., Smith, S.M., 2018. Image processing and Quality Control for the first 10,000 brain imaging datasets from UK Biobank. Neuroimage 166, 400–424. https://doi.org/10.1016/j.neuroimage.2017.10.034

Brown, T.T., Kuperman, J.M., Chung, Y., Erhart, M., McCabe, C., Hagler, D.J., Venkatraman, V.K., Akshoomoff, N., Amaral, D.G., Bloss, C.S., Casey, B.J., Chang, L., Ernst, T.M., Frazier, J.A., Gruen, J.R., Kaufmann, W.E., Kenet, T., Kennedy, D.N., Murray, S.S., Sowell, E.R., Jernigan, T.L., Dale, A.M., 2012. Neuroanatomical assessment of biological maturity. Curr. Biol. 22, 1693–1698. https://doi.org/10.1016/j.cub.2012.07.002

Bzdok, D., Varoquaux, G., Steyerberg, E.W., 2020. Prediction, not association, paves the road to precision medicine. JAMA Psychiatry. https://doi.org/10.1001/jamapsychiatry.2020.2549

Cole, J., Raffel, J., Friede, T., Eshaghi, A., Brownlee, W., Chard, D., De Stefano, N., Enzinger, C., Pirpamer, L., Filippi, M., Gasperini, C., Rocca, M., Rovira, A., Ruggieri, S., Sastre-Garriga, J., Stromillo, M., Uitdehaag, B., Vrenken, H., Barkhof, F., Nicholas, R., Ciccarelli, O., 2019. Accelerated brain ageing and disability in multiple sclerosis. bioRxiv 584888. https://doi.org/10.1101/584888

Cole, J.H., Franke, K., 2017. Predicting Age Using Neuroimaging: Innovative Brain Ageing Biomarkers. Trends Neurosci. 40, 681–690. https://doi.org/10.1016/j.tins.2017.10.001

Cole, J.H., Poudel, R.P.K., Tsagkrasoulis, D., Caan, M.W.A., Steves, C., Spector, T.D., Montana, G., 2017. NeuroImage Predicting brain age with deep learning from raw imaging data results in a reliable and heritable biomarker. Neuroimage 163, 115–124. https://doi.org/10.1016/j.neuroimage.2017.07.059

Cole, J.H., Ritchie, S.J., Bastin, M.E., Valdés Hernández, M.C., Muñoz Maniega, S., Royle, N., Corley, J., Pattie, A., Harris, S.E., Zhang, Q., Wray, N.R., Redmond, P., Marioni, R.E., Starr, J.M., Cox, S.R., Wardlaw, J.M., Sharp, D.J., Deary, I.J., 2018. Brain age predicts mortality. Mol. Psychiatry 23, 1385–1392. https://doi.org/10.1038/mp.2017.62

Dinsdale, N.K., Jenkinson, M., Namburete, A.I.L., 2020. Deep Learning-Based Unlearning of Dataset Bias for MRI Harmonisation and Confound Removal. bioRxiv 2020.10.09.332973. https://doi.org/10.1101/2020.10.09.332973

Dosenbach, N.U.F., Nardos, B., Cohen, A.L., Fair, D.A., Power, J.D., Church, J. a, Nelson, S.M., Wig, G.S., Vogel, A.C., Coalson, R.S., Jr, J.R.P., Barch, D.M., 2010. Prediction of Individual Brain Maturity Using fMRI. Science (80-.). 329, 1358–1361. http://dx.doi.org/10.1016/j.jrurstud.2015.06.009

Franke, K., Ziegler, G., Klöppel, S., Gaser, C., 2010. Estimating the age of healthy subjects from T1-weighted MRI scans using kernel methods: Exploring the influence of various parameters. Neuroimage 50, 883–892. https://doi.org/10.1016/j.neuroimage.2010.01.005

Kaufmann, T., van der Meer, D., Doan, N.T., Schwarz, E., Lund, M.J., Agartz, I., Alnæs, D., Barch, D.M., Baur-Streubel, R., Bertolino, A., Bettella, F., Beyer, M.K., Bøen, E., Borgwardt, S., Brandt, C.L., Buitelaar, J., Celius, E.G., Cervenka, S., Conzelmann, A., Córdova-Palomera, A., Dale, A.M., de Quervain, D.J.F., Carlo, P., Djurovic, S., Dørum, E.S., Eisenacher, S., Elvsåshagen, T., Espeseth, T., Fatouros-Bergman, H., Flyckt, L., Franke, B., Frei, O., Haatveit, B., Håberg, A.K., Harbo, H.F., Hartman, C.A., Heslenfeld, D., Hoekstra, P.J., Høgestøl, E.A., Jernigan, T.L., Jonassen, R., Jönsson, E.G., Kirsch, P., Kłoszewska, I., Kolskår, K.K., Landrø, N.I., Hellard, S., Lesch, K.-P., Lovestone, S., Lundervold, A., Lundervold, A.J., Maglanoc, L.A., Malt, U.F., Mecocci, P., Melle, I., Meyer-Lindenberg, A., Moberget, T., Norbom, L.B., Nordvik, J.E., Nyberg, L., Oosterlaan, J., Papalino, M., Papassotiropoulos, A., Pauli, P., Pergola, G., Persson, K., Richard, G., Rokicki, J., Sanders, A.-M., Selbæk, G., Shadrin, A.A., Smeland, O.B., Soininen, H., Sowa, P., Steen, V.M., Tsolaki, M., Ulrichsen, K.M., Vellas, B., Wang, L., Westman, E., Ziegler, G.C., Zink, M., Andreassen, O.A., Westlye, L.T., 2019. Common brain disorders are associated with heritable patterns of apparent aging of the brain. Nat. Neurosci. 22, 1617–1623. https://doi.org/10.1038/s41593-019-0471-7

Kingma, D.P., Ba, J.L., 2014. Adam: A method for stochastic optimization. arXiv Prepr. arXiv1412.6980.

Lanka, P., Rangaprakash, D., Dretsch, M.N., Katz, J.S., Denney, T.S., Deshpande, G., 2019. Supervised machine learning for diagnostic classification from large-scale neuroimaging datasets. Brain Imaging Behav. 1–39. https://doi.org/10.1007/s11682-019-00191-8

LeCun, Y., Bengio, Y., Hinton, G., 2015. Deep learning. Nature 521, 436–444. https://doi.org/10.1038/nature14539

Levakov, G., Rosenthal, G., Shelef, I., Raviv, T.R., Avidan, G., 2020. From a deep learning model back to the brain—Identifying regional predictors and their relation to aging. Hum. Brain Mapp. hbm.25011. https://doi.org/10.1002/hbm.25011

Liu, M., Zhang, J., Adeli, E., Shen, D., 2018. Landmark-based deep multi-instance learning for brain disease diagnosis. Med. Image Anal. 43, 157–168. https://doi.org/10.1016/j.media.2017.10.005

Miller, K.L., Alfaro-Almagro, F., Bangerter, N.K., Thomas, D.L., Yacoub, E., Xu, J., Bartsch, A.J., Jbabdi, S., Sotiropoulos, S.N., Andersson, J.L.R., Griffanti, L., Douaud, G., Okell, T.W., Weale, P., Dragonu, I., Garratt, S., Hudson, S., Collins, R., Jenkinson, M., Matthews, P.M., Smith, S.M., 2016. Multimodal population brain imaging in the UK Biobank prospective epidemiological study. Nat. Neurosci. 19, 1523–1536. https://doi.org/10.1038/nn.4393

Neeb, H., Zilles, K., Shah, N.J., 2006. Fully-automated detection of cerebral water content changes: Study of age-and gender-related H2O patterns with quantitative MRI. Neuroimage 29, 910–922. https://doi.org/10.1016/j.neuroimage.2005.08.062

Peng, H., Gong, W., Beckmann, C.F., Vedaldi, A., Smith, S.M., 2019. Accurate brain age prediction with lightweight deep neural networks. bioRxiv879346.

Raghu, M., Zhang, C., Kleinberg, J., Bengio, S., 2019. Transfusion: Understanding Transfer Learning for Medical Imaging, in: Advances in Neural Information Processing Systems 32. Curran Associates, Inc., pp. 3347–3357.

Rosenberg, M.D., Casey, B.J., Holmes, A.J., 2018. Prediction complements explanation in understanding the developing brain. Nat. Commun. 9, 1–13. https://doi.org/10.1038/s41467-018-02887-9

Russakovsky, O., Deng, J., Su, H., Krause, J., Satheesh, S., Ma, S., Huang, Z., Karpathy, A., Khosla, A., Bernstein, M., Berg, A.C., Fei-Fei, L., 2015. ImageNet Large Scale Visual Recognition Challenge. Int. J. Comput. Vis. 115, 211–252. https://doi.org/10.1007/s11263-015-0816-y

Scheinost, D., Noble, S., Horien, C., Greene, A.S., Lake, E.M., Salehi, M., Gao, S., Shen, X., O’Connor, D., Barron, D.S., Yip, S.W., Rosenberg, M.D., Constable, R.T., 2019. Ten simple rules for predictive modeling of individual differences in neuroimaging. Neuroimage. https://doi.org/10.1016/j.neuroimage.2019.02.057

Shmueli, G., 2010. To explain or to predict? Stat. Sci. 25, 289–310. https://doi.org/10.1214/10-STS330

Smith, S.M., Douaud, G., Chen, W., Hanayik, T., Alfaro-Almagro, F., Sharp, K., Elliott, L.T., 2020a. Enhanced Brain Imaging Genetics in UK Biobank. bioRxiv 2020.07.27.223545. https://doi.org/10.1101/2020.07.27.223545

Smith, S.M., Elliott, L.T., Alfaro-Almagro, F., McCarthy, P., Nichols, T.E., Douaud, G., Miller, K.L., 2020b. Brain aging comprises many modes of structural and functional change with distinct genetic and biophysical associations. Elife 9, 1–28. https://doi.org/10.7554/eLife.52677

Smith, S.M., Jenkinson, M., Woolrich, M.W., Beckmann, C.F., Behrens, T.E.J., Johansen-Berg, H., Bannister, P.R., De Luca, M., Drobnjak, I., Flitney, D.E., Niazy, R.K., Saunders, J., Vickers, J., Zhang, Y., De Stefano, N., Brady, J.M., Matthews, P.M., 2004. Advances in functional and structural MR image analysis and implementation as FSL, in: NeuroImage. Academic Press, pp. S208–S219. https://doi.org/10.1016/j.neuroimage.2004.07.051

Smith, S.M., Vidaurre, D., Alfaro-Almagro, F., Nichols, T.E., Miller, K.L., 2019. Estimation of brain age delta from brain imaging. Neuroimage 200, 528–539. https://doi.org/10.1016/j.neuroimage.2019.06.017

Thomas, R.M., Gallo, S., Cerliani, L., Zhutovsky, P., El-Gazzar, A., van Wingen, G., 2020. Classifying Autism Spectrum Disorder Using the Temporal Statistics of Resting-State Functional MRI Data With 3D Convolutional Neural Networks. Front. Psychiatry 11, 1. https://doi.org/10.3389/fpsyt.2020.00440

Zeng, L.L., Wang, H., Hu, P., Yang, B., Pu, W., Shen, H., Chen, X., Liu, Z., Yin, H., Tan, Q., Wang, K., Hu, D., 2018. Multi-Site Diagnostic Classification of Schizophrenia Using Discriminant Deep Learning with Functional Connectivity MRI. EBioMedicine 30, 74–85. https://doi.org/10.1016/j.ebiom.2018.03.017

Zou, L., Zheng, J., Miao, C., McKeown, M.J., Wang, Z.J., 2017. 3D CNN Based Automatic Diagnosis of Attention Deficit Hyperactivity Disorder Using Functional and Structural MRI. IEEE Access 5, 23626–23636. https://doi.org/10.1109/ACCESS.2017.2762703

